# Metabolic switching is impaired by aging and facilitated by ketosis independent of glycogen

**DOI:** 10.1101/2019.12.12.874297

**Authors:** Abbi R. Hernandez, Leah M. Truckenbrod, Quinten P. Federico, Keila T. Campos, Brianna M. Moon, Nedi Ferekides, Meagan Hoppe, Dominic D’Agostino, Sara N. Burke

## Abstract

The ability to switch between glycolysis and ketosis promotes survival by enabling metabolism through fat oxidation during periods of fasting. Carbohydrate restriction or stress can also elicit metabolic switching. Keto-adapting from glycolysis is delayed in aged rats, but factors mediating this age-related impairment have not been identified. We measured metabolic switching between glycolysis and ketosis, as well as glycogen dynamics, in young and aged rats undergoing time-restricted feeding (TRF) with a standard diet or a low carbohydrate ketogenic diet (KD). TRF alone reversed markers of insulin-related metabolic deficits and accelerated metabolic switching in aged animals. A KD+TRF, however, provided additive benefits on these variables. Remarkably, the ability to keto-adapt was not related to glycogen levels and KD-fed rats showed an enhanced elevation in glucose following epinephrine administration. This study provides new insights into the mechanisms of keto-adaptation demonstrating the utility of dietary interventions to treat metabolic impairments across the lifespan.

## INTRODUCTION

Patterns of energy consumption are a powerful regulator of mortality and longevity. In addition to caloric restriction, which has been extensively shown to extend maximum lifespan across species (de Cabo et al., 2014; Fontana, 2007; Fontana et al., 2010; Longo and Mattson, 2014; Longo and Panda, 2016; Masoro, 2005; Mattson et al., 2014), time restricted feeding (TRF; also referred to as meal feeding or intermittent fasting) may also improve morbidity and mortality rates (Mitchell et al., 2019). Intermittent periods of eating inevitably lead to episodes of ketogenesis, in which fat is oxidized into ketone bodies, reducing the need for glycolysis (Anson et al., 2003; Mitchell et al., 2019, 2016).

Without fasting, ketosis can also be achieved with high fat, low carbohydrate ketogenic diets (KD). Importantly, long-term ketosis has also been shown to prevent cognitive and physical declines associated with aging (Bredesen, 2014; Choi et al., 2018; Greco et al., 2016; Hernandez et al., 2018a; Hernandez et al., 2018; Hussain et al., 2012; Kephart et al., 2017; Westman et al., 2008). It remains to be determined, however, whether TRF alone can produce sufficient levels of ketosis to elicit similar health benefits and the extent to which TRF with a KD may have additive benefits on the healthspan of older adults and other animals.

The process of switching from glycolysis to a reliance on fat-derived ketone bodies as a primary energy source is termed keto-adaptation (Phinney, 2004; Volek et al., 2016). While nutritional ketosis has been proposed as an intervention for a number of age-related metabolic and neurological diseases (Gasior et al., 2006; Hartman, 2012; Paoli et al., 2013), aged animals take longer than young to produce high levels of circulating ketone bodies (Hernandez et al., 2018; Hernandez et al., 2018a). Understanding the factors responsible for delayed keto-adaptation in old animals, and how to overcome this, are critical as brain utilization of ketones bodies directly correlates with circulating levels in the periphery (Bentourkia et al., 2009; Daniel et al., 1971; Hawkins et al., 1986). Thus, in order to design precision ketogenic diet therapy (Lukosaityte et al., 2017) that can improve cognitive functioning and physical health in older adults, it is imperative to understand ketogenesis across the lifespan and the metabolic variables that can delay or promote metabolic switching.

The prevailing theory regarding the initiation of ketogenesis is that ketone body production is triggered once the body depletes its glycogen stores. This glycogen depletion hypothesis likely stems from the body’s natural metabolic response to vigorous exercise or longer fasts (Dohm et al., 1983; Ivy, 1991). Following 1 hour of exercise, there is an inverse relationship between glycogen content and ketone body levels (Adams and Koeslag, 1988). Given the presumed relationship between keto-adaptation and glycogen levels, it is conceivable that aged rats take longer to achieve stable nutritional ketosis due to higher levels of glycogen or altered glycogen dynamics. There are differences in body mass with age (Hernandez et al., 2018), as well as evidence that aged rats have reduced glycogenolytic capacity (Morris et al., 2010). Specifically, the hepatic release of glycogen (Talley et al., 2000), which increases blood glucose in response to epinephrine (McNay and Gold, 2001), is decreased in aged compared to young rats (Morris and Gold, 2013). Thus, old animals may be deficient at converting glycogen to glucose on demand, which would decrease the rate of glycogen depletion. An alternative to the glycogen depletion hypothesis is that old animals have impaired metabolic switching, herein defined as the ability to utilize either carbohydrates or fat as a fuel source when appropriate. In line with this idea, insulin insensitivity occurs in advanced age (Fink et al., 1983; Refaie et al., 2006; Ryan, 2000), and fasting insulin levels are often elevated. Since insulin inhibits fat oxidation, higher insulin levels could also delay keto-adaptation. The current study tested the extent to which altered keto-adaptation in old age was related to impaired glycogen dynamics or metabolic dysfunction in rats that were placed on TRF (fed once per day) with a standard diet (SD) or TRF in combination with an isocaloric ketogenic diet (KD). For some comparisons, a third group of rats fed *ad libitum* (also referred to as free feeding) standard rodent chow was included.

## RESULTS

### Age-related delays in keto-adaptation are not due to altered feeding patterns

Keto-adaptation is the coordinated set of metabolic adaptations that ensures proper interorgan fuel supply in the face of low carbohydrate availability (Volek et al., 2015). This process is enabled by metabolic switching from glycolysis to ketosis in which glucose oxidation is replaced by lipid oxidation and the synthesis of ketone bodies (primarily beta-hydroxybutyrate and acetoacetate) in the liver. A proxy for keto-adaptation is elevated, asymptotic levels of blood beta-hydroxybutyrate in response to consumption of a high-fat, low carbohydrate ketogenic diet (KD).

A potential explanation for delayed keto-adaptation, as previously observed in aged compared to young rats (Hernandez et al., 2019a, 2018), could be altered food consumption patterns. To rule out differences in food consumption, and to evaluate the amount of time that rats were fasted during TRF, caloric intake was measured for three hours postprandially and at 24 hours. Young (4 mo) and aged (20 mo) male Fischer 344 × Brown Norway F1 Hybrid rats fed 51 kcal of the SD or KD once per day, as well as young and aged rats free-fed regular chow *ad libitum*, were evaluated for this comparison. The *ab libitum* fed groups allowed for a comparison of food consumption patterns during TRF, as used here, and *ad libitum* feeding that is common in aging studies.

ANOVA-RM with the between subjects factors of age and diet group indicated a significant effect of time point (F_[8,224]_ = 329.96; p < 0.001; Figure 1C) on the amount of food consumed for all groups, as well as a significant effect of diet group (F_[2,28]_ = 35.86; p < 0.001), but no effect of age (F_[1,28]_ = 0.58; p = 0.45). Critically, the consumption patterns of young and aged rats on the SD or KD did not vary, indicating that altered caloric intake over time cannot account for the age-related delay in keto-adaptation. Diet group, however, significantly interacted with time point (F_[16,224]_ = 67.59; p < 0.001). Specifically, the *ad libitum*-fed rats did not consume the same density of calories in the 3-hour postprandial period, and consumed significantly more calories 24-hr after feeding (F_[3,30]_ = 113.326; p < 0.001). Furthermore, after 24 hours, the aged *ad libitum*-fed rats consumed significantly more kcal than the young rats of the same feeding group.

**Figure 1:**
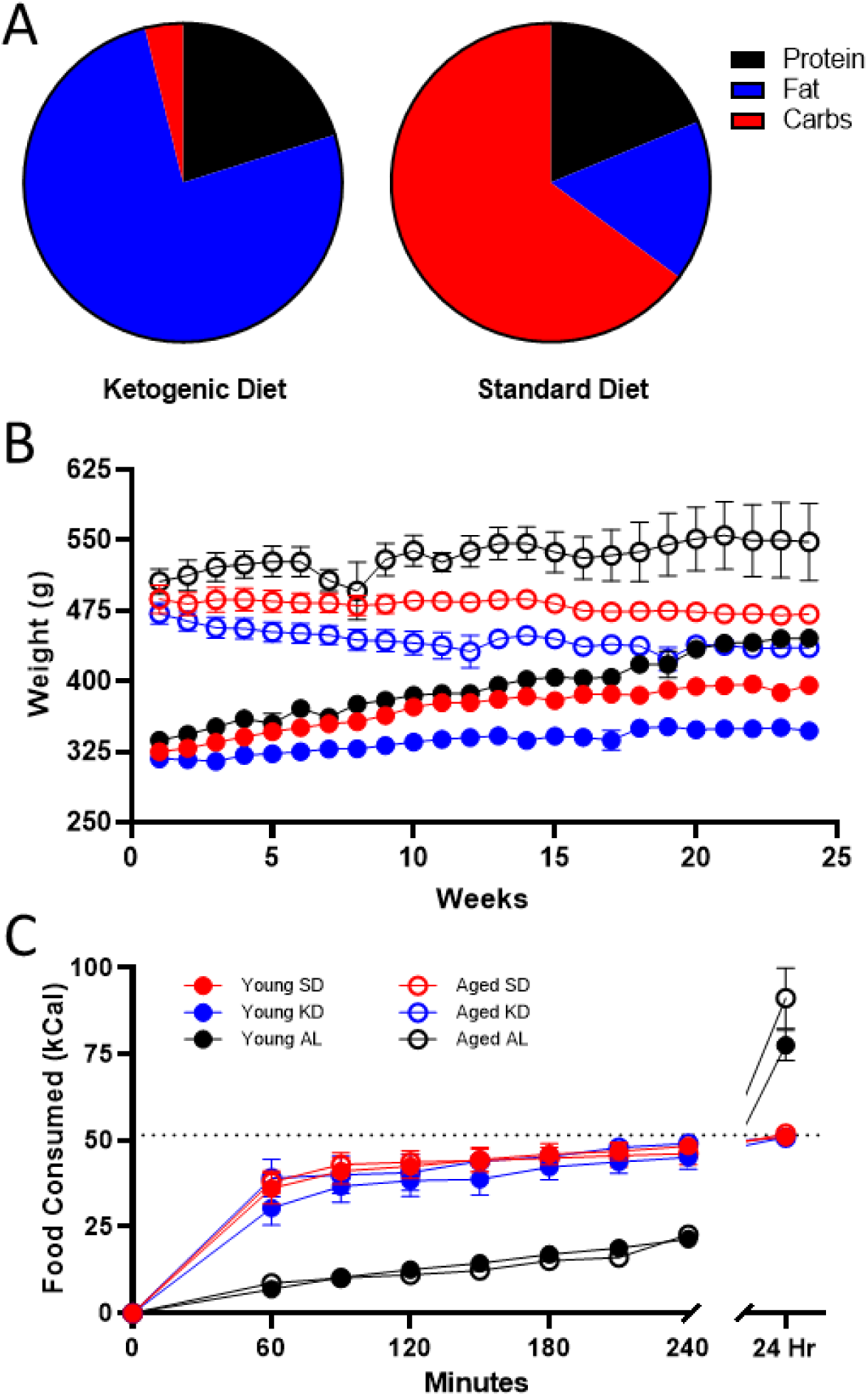
Dietary implementation. (A) Ketogenic (KD) and standard diet (SD) macronutrient compositions. (B) Weight significantly differed across age groups throughout the study, but most notably was significantly different in *ad libitum* fed rats relative to time restricted fed (TRF) rats. (C) The number of kcal significantly differed across the two feeding methods, as *ad libitum*-fed rats consumed at a slower rate than TRF rats, but consumed significantly more during a 24 hour period. Dotted line indicates kcal allotment given to TRF groups (∼51 kcal). Data are represented as group mean ±1 standard error of the mean (SEM).

Because the caloric intake in the SD and KD groups was significantly less in a 24-hr period than animals that were fed *ad libitum*, we compared the body weights of these animals over the course of 6 months. Prior to diet implementation, there were no differences in weight across eventual diet groups (F_[2,32]_ = 0.33; p = 0.72), though aged rats weighed significantly more than young (F_[1,32]_ = 588.96; p < 0.001). ANOVA-RM indicated a significant main effect of week (F_[23,644]_ = 12.89; p < 0.001), age (F_[1,28]_ = 250.67; p < 0.001) and diet (F_[2,28]_ = 27.16; p < 0.001) across 24 weeks. Furthermore, both age (F_[23,644]_ = 17.72; p < 0.001) and diet (F_[23,644]_ = 6.38; p < 0.001) significantly interacted with week number. However, there was no significant interaction between age and diet (F_[2,28]_ = 1.45; p = 0.25). This pattern in the data is due, at least in part, to the *ad libitum*-fed animals gaining significantly more weight than TRF rats. In fact, *ad libitum*-fed rats had significant weight gain across the 6 months for both young (F_[1,18]_ = 29.35; p < 0.001) and aged rats (F_[1,18]_ = 8.71; p < 0.02). It should be noted, however, that even on TRF of 51 kcal/day, both SD (t_[6]_ = 8.60; p < 0.001) and KD (t_[6]_ = 4.49; p < 0.01) fed young mature adult rats weighed significantly more at the end of the 6 months. These data demonstrate that pathological overconsumption was prevented in the current animals without caloric restriction.

After establishing that food consumption patterns did not vary between age groups, postprandial blood glucose (Figure 2A) and ketone body levels (BHB; Figure 2B) during the first day of initiation of a medium chain triglyceride (MCT) oil-based KD or a calorically-matched and micronutrient equivalent standard diet (SD). This procedure was done to examine if age-related delays in keto-adaptation could be replicated in these animals. Blood measurements were conducted every 3 days for 28 days in rats that underwent TRF with access to 51 kcal given once daily. In both young and aged rats, the KD decreased blood glucose levels and increased serum BHB. The reduction in glucose and concurrent increase in BHB can be combined to quantify the overall level of ketosis using the glucose ketone index (GKI; Figure 2C). Importantly, the GKI tracks better with therapeutic efficacy of KDs in cancer than either glucose or BHB levels alone (Meidenbauer et al., 2015). Briefly, the GKI is the ratio of the molecular weight normalized serum glucose level (expressed as mol/g × dL), to the serum BHB levels (mM). Thus, lower GKI values are indicative of higher levels of ketosis. Therefore, the time required for GKI to be asymptotically lowered can be used to extrapolate the time it takes an animal to keto-adapt.

**Figure 2:**
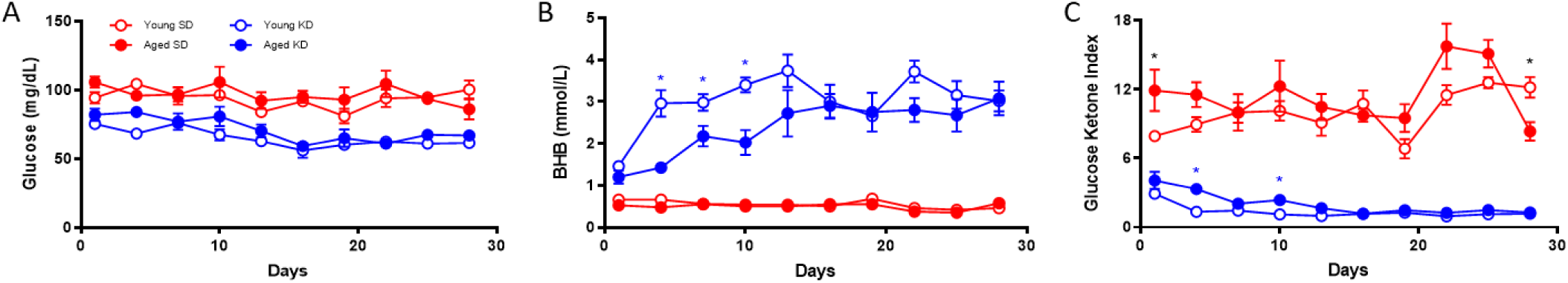
Keto-adaptation differs by age. (A) Glucose, (B) BHB and (C) GKI levels of young and aged rats consuming either a KD or SD as a function of days on the diet indicate an age-dependent delay in keto-adaptation. Data are represented as group mean ±1 standard error of the mean (SEM), *p<0.05.

During 1 month of TRF with either the isocaloric KD or SD, the GKI was significantly lower in KD-fed animals (F_[1,14]_ = 1459.55; p < 0.001). Moreover, the GKI was significantly lower in young compared to aged rats (F_[1,14]_ = 20.23; p = 0.001), indicating lower levels of ketosis in older animals. When examined longitudinally, the GKI significantly changed across the 28 days (F_[9,126]_ = 4.21; p < 0.001). The interaction between days on the diet and diet group was also significant (F_[9,126]_ = 6.12; p < 0.001), such that the GKI in KD-fed rats significantly decreased while the GKI of SD-fed rats did not. Importantly, GKI levels across days on the diet significantly interacted with age (F_[9,126]_ = 2.78; p < 0.01), with young rats having significantly lower GKI values compared to aged rats on days 4 (t_[7]_ = 7.11; p = 0.001) and 10 (t_[7]_ = 3.14; p = 0.02). No difference across age groups was detected from day 13 onward (p > 0.1). These data replicate a previous observation that aged rats require longer than young to keto-adapt, but can maintain stable levels of ketogenesis thereafter (Hernandez et al., 2018).

### Age-related metabolic impairments delay keto-adaptation rate

To assess the contribution of age-related metabolic differences on keto-adaptation, young and aged rats fed *ad libitum* underwent an insulin tolerance test (ITT). Insulin powerfully inhibits ketogenesis (Fukao et al., 2004), therefore insulin receptor insensitivity and hyperinsulinemia could delay keto-adaptation. Notably, all animals were fed *ad libitum* prior to initiating the KD or SD, so these rats represent the basal metabolic state prior to experimental diets being implemented. ANOVA-RM with the between subjects factor of age indicated a significant effect of time point (F_[6,54]_ = 83.33; p < 0.001), but glucose levels at baseline were not significantly different between age groups (t_[9]_ = 1.70; p = 0.12; Figure 3A). The area under the curve for glucose (Figure 3B/C) was significantly greater for aged rats relative to young (t_[9]_ = 3.55; p = 0.006), demonstrating an impaired glucose response to the ITT. BHB levels also did not significantly differ at baseline across age groups (t_[9]_ = 1.48; p = 0.17; Figure 3D), but BHB declined in response to the insulin injection (Figure 3E). However, the area under the curve did not differ across age groups (t_[9]_ = 0.38; p = 0.71; Figure 3E/F).

**Figure 3:**
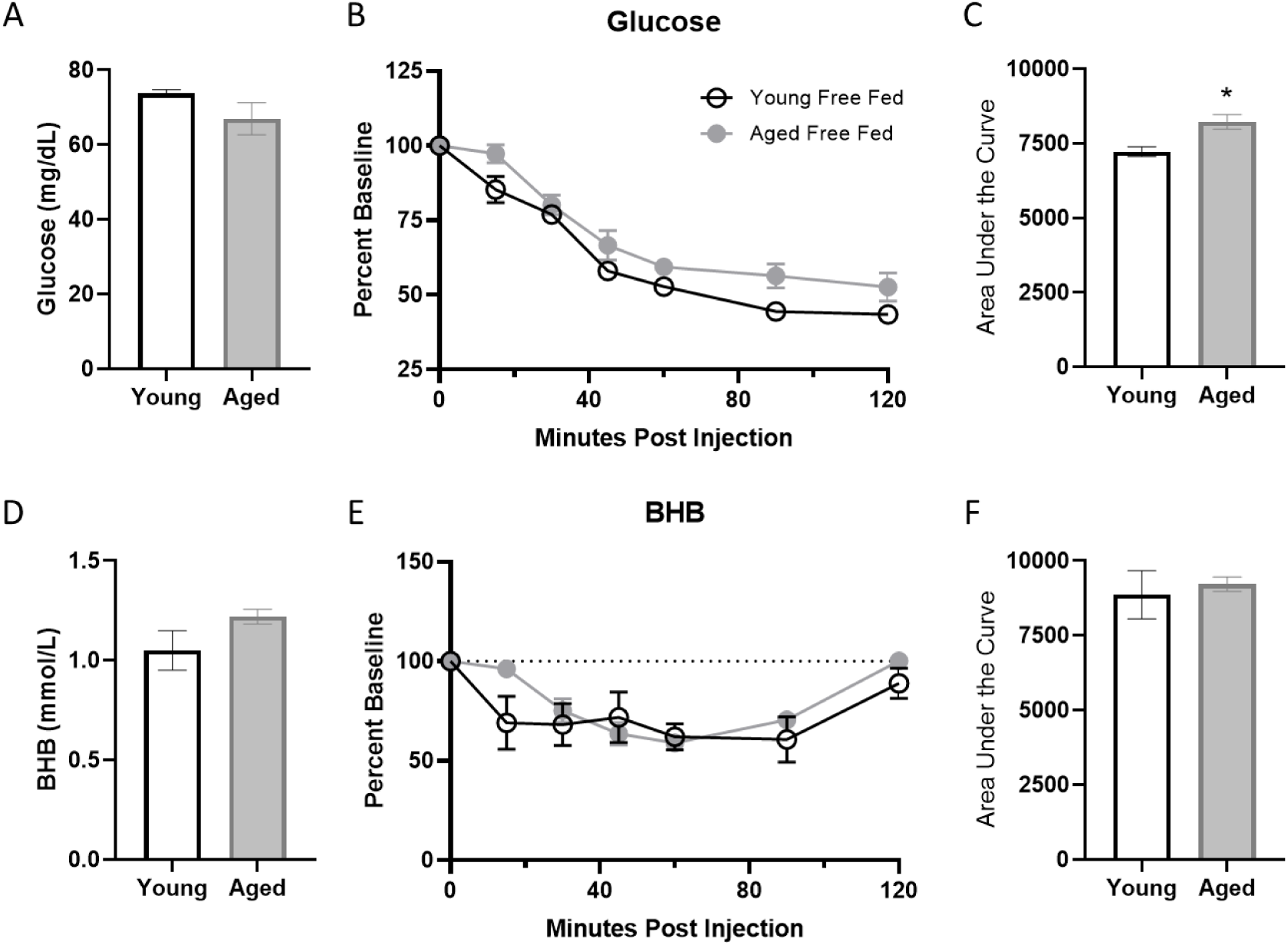
Insulin tolerance is impaired in aged *ad libitum-*fed rats. Glucose response (A) was not different at baseline (time 0) across age groups, but both the response to insulin expressed as (B) percent of baseline over time and (C) the area under the curve (AUC) was significantly elevated in aged rats relative to young. However, there were no age differences in β-hydroxybutyrate (BHB) (D) at baseline, (E) over time following insulin injection or (F) in total AUC in *ad libitum*-fed rats. Data are represented as group mean ±1 SEM. * = p < 0.05.

To further characterize metabolic impairments between young and aged *ad libitum*-fed rats, levels of several markers of peripheral metabolism were obtained (Figure 4). When fasted, aged rats had significantly higher insulin (t_[29]_ = 3.14; p = 0.004), leptin (t_[49]_ = 3.54; p < 0.001) and C-peptide (t_[48]_ = 3.41; p < 0.01). In a subset of these rats, fasted glucose was also obtained and a HOMA value could be calculated as (insulin (mU/I) × glucose (mg/dL)) / 405. Aged rats had a significant elevation in HOMA relative to young rats (t_[20]_ = 3.36; p = 0.003; Figure 4D). Together these data show that aged rats fed *ad libitum* have metabolic impairments and it is notable that elevated HOMA values are a risk factor for diabetes type II (Meigs et al., 2007) as well as cardiovascular disease (Bonora et al., 2002). Because insulin inhibits ketosis, the hyperinsulinemia in aged rats could be responsible for delayed keto-adaptation in aging.

**Figure 4:**
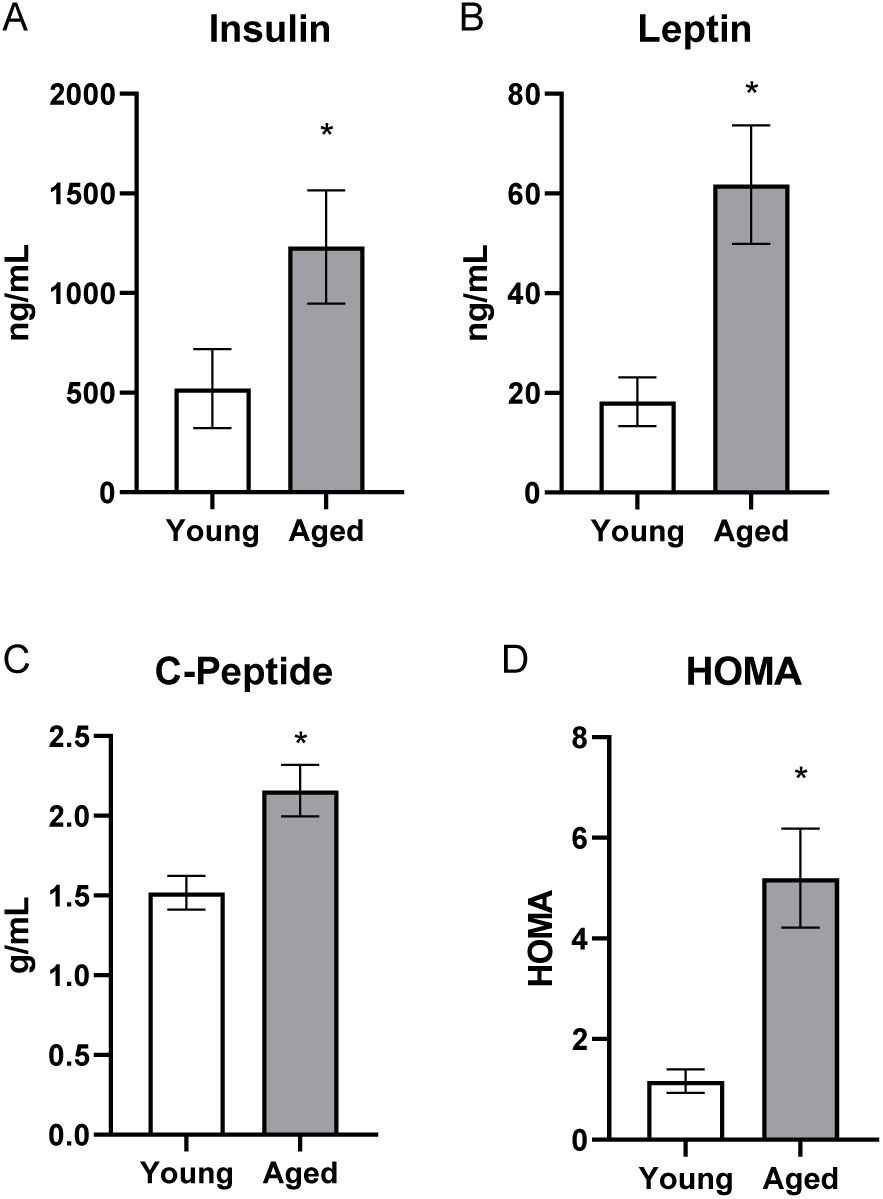
Aged rats have impaired metabolic parameters when fed *ad libitum*. Aged rats had significantly elevated (A) insulin, (B) leptin, (C) C-peptide and (D) HOMA levels relative to young rats when fed standard rodent chow *ad libitum*. Data are represented as group mean ±1 SEM. * = p < 0.05.

Because aged rats allowed to eat *ad libitum* had significant impairments in insulin regulation, the potential of both TRF with the SD and TRF with the KD to reverse insulin impairments was investigated during a 4-week period of keto-adaptation. Fasted insulin, leptin, and C-peptide levels, as well as glucose levels for the generation of HOMA values, were measured in young and aged rats undergoing TRF with either the SD or KD for 1, 4, 7 or 28 days. Insulin inhibits ketogenesis (Fukao et al., 2004), therefore, higher basal levels of insulin in aged rats could account for delayed keto-adaptation.

Fasted insulin levels differed significantly across diet groups (F_[1,64]_ = 17.74; p < 0.001), but not age groups (F_[1,64]_ = 1.64; p = 0.21). However, age and diet did significantly interact (F_[1,64]_ = 7.75; p = < 0.01). While there was no overall main effect of length of time on the diet (F_[3,64]_ = 0.40; p = 0.76), length of time on the diet also significantly interacted with age (F_[3,64]_ = 4.69; p < 0.01) but not diet group (F_[3,64]_ = 0.45; p = 0.72). In animals on the KD, fasted insulin levels were significantly greater in aged compared to young animals (F_[1,32]_ = 13.35; p = 0.001; Figure 5A). Interestingly, this age difference was not evident on day 28 of the KD (t_[8]_ = 0.28; p = 0.79). There was no difference across age (F_[1,32]_ = 0.82; p = 0.37) in SD-fed rats over 28 days of TRF, however aged rats had significantly lower fasted insulin than young at day 28 (t_[8]_ = 2.49; p = 0.04; Figure 5B).

**Figure 5:**
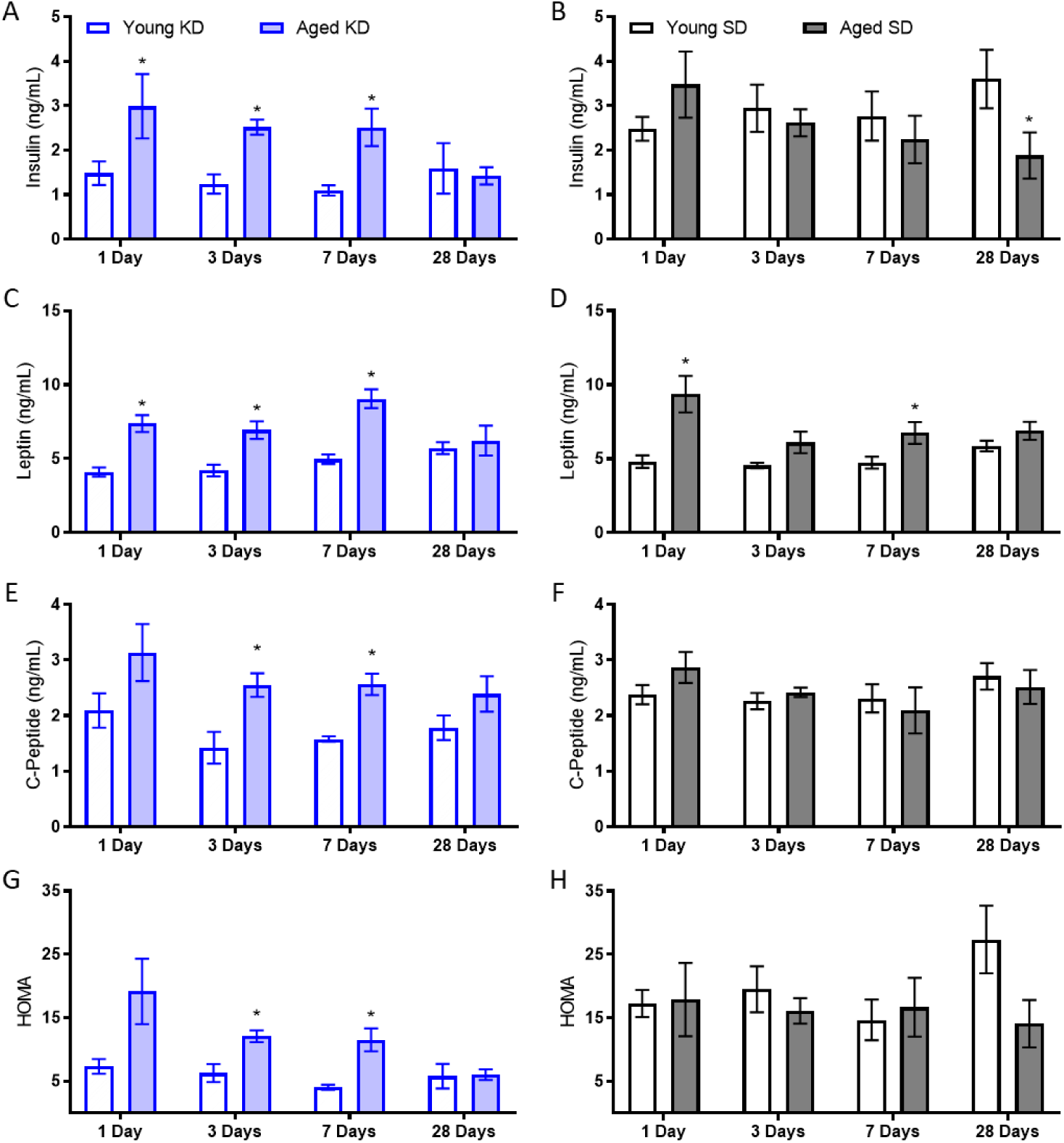
Insulin levels while keto-adapting. (A) Young rats fed a KD demonstrated immediate decreases in insulin levels. Aged rats did not demonstrate similar declines until after the first week of adaptation. (B) Neither young nor aged rats fed the SD, even in conjunction with TRF, demonstrated changes in insulin levels until the end of the 4^th^ week, at which point aged rats demonstrated decreased insulin levels. While aged rats on both (C) the KD and (D) the SD had significantly higher leptin levels than young rats on the same diet, only aged KD-fed rats demonstrated significant decrease over time. (D) C-Peptide levels were significantly lower in young rats relative to aged when fed a KD (E) but not an SD, though neither group demonstrated significant differences over time. (G) HOMA values were significantly higher in aged KD-fed rats relative to young at the start of the diet, but this difference was ameliorated by day 28. (H) However, aged rats fed an SD did not see improvements in HOMA over time. Data are represented as group mean ±1 SEM. * = p < 0.05.

Leptin was significantly elevated in aged rats relative to young (F_[1,64]_ = 64.21; p < 0.001), but was not altered by diet or time groups (p > 0.1 for both). However, time group significantly interacted with both age (F_[3,64]_ = 4.76; p < 0.01) and diet group (F_[3,64]_ = 3.19; p = 0.03). When diet groups were compared individually, leptin was significantly higher in aged rats fed both the KD (F_[1,32]_ = 42.99; p < 0.001) and SD (F_[1,32]_ = 24.15; p < 0.001). While time point did not reach significance for either diet group (KD p = 0.06; SD p = 0.07), there was a significant interaction between age and time point for KD fed rats only (F_[3,32]_ = 3.57; p = 0.03).

C-peptide levels were also significantly elevated in aged relative to young rats (F_[1,64]_ = 13.37; p < 0.01). While both trended towards significance, neither diet (F_[1,64]_ = 3.45; p = 0.07) nor time point (F_[3,64]_ = 2.64; p = 0.06) significantly affected C-peptide levels. However, the significant interaction between age and diet (F_[1,64]_ = 10.34; p < 0.01) indicates that age was a significant factor for the KD-fed rats but not SD-fed rats. Therefore, the 2 diet groups were considered individually. C-peptide did not change significantly over time for either SD (F_[3,32]_ = 1.32; p = 0.29) nor KD (F_[3,32]_ = 1.95; p = 0.14). However, C-peptide levels were significantly lower in young rats relative to aged rats on the KD (F_[1,32]_ = 20.96; p < 0.001) but not SD (F_[1,32]_ = 0.11; p = 0.74), though this age difference was not present at 28 days.

Lastly, glucose levels at the time of blood collection were combined with insulin levels to derive a HOMA value (see above). While age did not significantly affect HOMA values overall (F_[1,64]_ = 0.76; p = 0.39), age significantly interacted with diet group (F_[1,64]_ = 9.24; p < 0.01) and time group (F_[1,64]_ = 3.09; p = 0.03). Furthermore, the KD significantly lowered HOMA values (F_[1,64]_ = 30.33; p < 0.001), so diet groups were analyzed separately. Aged, KD-fed rats had significantly higher HOMA values than young (F_[1,32]_ = 16.54; p < 0.001), and this significantly changed over the 28-day keto-adaptation period (F_[1,32]_ = 4.00; p = 0.02) such that by day 28, aged rats did not differ from young (t_[8]_ = 0.11; p = 0.91). Conversely, aged, SD-fed rats did not exhibit higher HOMA values than young (F_[1,32]_ = 1.53; p = 0.23), and HOMA values were not affected by length of time across the 28 days of TRF (F_[1,32]_ = 0.54; p = 0.66).

To investigate the potential of TRF and the KD to reverse impaired insulin tolerance in aged subjects, rats were given an insulin tolerance test (ITT) before and after 8 weeks of dietary intervention. At baseline, while all rats were consuming chow *ad libitum*, the insulin injection significantly affected peripheral glucose levels over time for all groups (F_[6,150]_ = 49.81; p < 0.001; data not shown). There were no differences in glucose response to insulin across assigned diet groups (F_[2,25]_ = 0.41; p = 0.67). Interestingly, ketone body levels were also significantly changed over time following an injection of insulin (F_[6,66]_ = 23.26; p < 0.001) such that they appeared to increase as the glucose level decreased. There were no differences across designated diet groups prior to diet implementation (F_[1,31]_ = 1.26; p = 0.30) in BHB response to insulin.

Following 8 weeks on their respective diets, the significant main effect of time point on glucose response to insulin remained significant across all groups (F_[6,156]_ = 199.23; p < 0.001; Figure 6). After the 8 weeks of experimental diets, however, there was a significant diet group effect (F_[2,26]_ = 10.12; p = 0.001), but no significant effect of age (F_[1,26]_ = 2.04; p = 0.17). Furthermore, time point significantly interacted with diet group (F_[12,156]_ = 3.13; p = 0.001) but not age (F_[12,156]_ = 0.43; p = 0.86). Because of the significant effects of diet group, glucose and BHB results are plotted separately for each diet group in Figures 6A-F. Notably, the AUC only significantly differed across age groups in the *ad libitum-*fed rats (t_[9]_ = 3.55; p = 0.006) but not the SD (t_[12]_ = 1.06; p = 0.31) or KD groups (t_[11]_ = 0.32; p = 0.75). These data suggest that age-related impairments in insulin sensitivity were ameliorated in both TRF groups.

**Figure 6:**
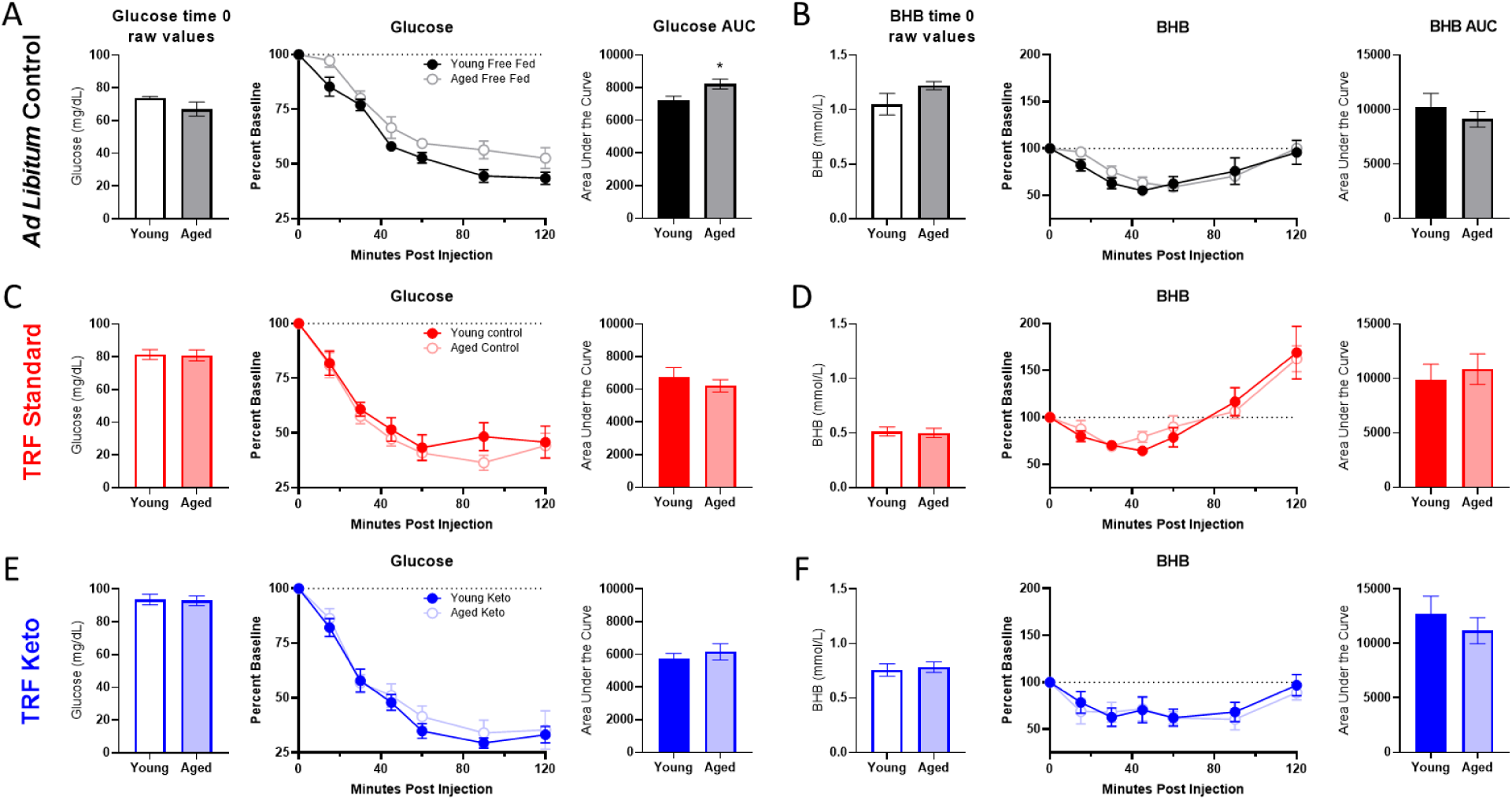
Insulin tolerance test (ITT) following 8 weeks of dietary implementation. (A) Replicating figure 4, at the 8 week time point there was a significant increase in the area under the curve for glucose in aged rats relative to young while fed *ad libitum* chow. (B) No differences in BHB were observed. (C-D) Neither glucose nor BHB values significantly differed across young and aged rats on the SD (E-F) nor the KD in response to a bolus of insulin injected intraperitoneally. Data are represented as group mean ±1 SEM. * p = < 0.05.

A glucose tolerance test (GTT) was also administered following 7 weeks of the diets. While the glucose injection did result in increased circulating glucose over time (F_[6,186]_ = 34.09; p < 0.001; Figure S1A), there were no differences across age (F_[1,31]_ = 0.14; p = 0.71) or diet groups (F_[2,31]_ = 1,15; p = 0.33), and none of these variables significantly interacted (p ≥ 0.10 for all comparisons). See supplementary material and Figure S1 for more information.

### Previous ketogenic diet experience enhances keto-adaptation beyond time-restricted feeding alone

The TRF implemented in the current experiments was associated with a ∼21 hour fast, as both young and aged rats consumed close to 100% of the available calories within 3 hours (Figure 1C). Because intermittent fasting (Mattson and Wan, 2005) and the KD (Ballard et al., 2013) have both been reported to improve insulin receptor sensitivity, as well as other markers of metabolic function, it is possible that either manipulation could enhance keto-adaptation in young and aged rats. To test this, young and aged rats were divided equally into groups A and B, with both groups receiving once daily TRF of 51 kcal as described previously with either the KD or SD for 6 weeks. Group A was placed on the KD for 4 consecutive weeks and then cycled onto the SD for 1 week, followed by a final week of the KD. Group B was placed on the SD for 4 weeks, followed by 1 week of the KD and a final week of the SD (Figure 7A). Blood glucose and BHB levels were collected every 3 days to calculate the GKI across the different diet cycles. Figure 7B shows the GKI values for groups A and B during KD initiation across the different cycles. During the cycles on the SD, all GKI values rapidly returned to high baseline levels on the first day and this did not vary between age groups (F_[1,14]_ = 1.35; p = 0.26; Figure 7C).

**Figure 7:**
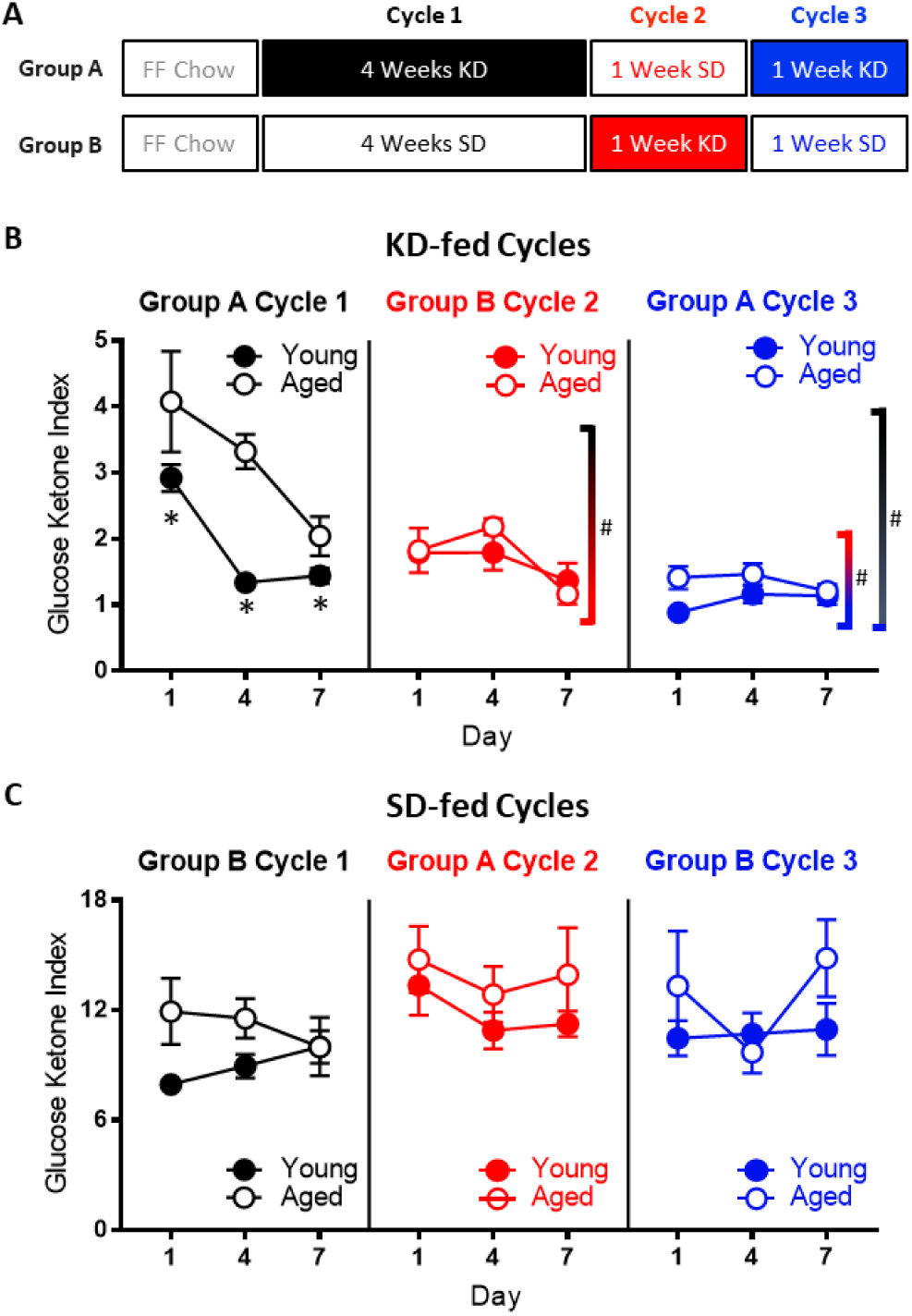
Age-related impairments in keto-adaptation are ameliorated with TRF and a previous keto-adaptation cycle. (A) Timeline of dietary interventions used for groups A and B. (B) GKI during 1 week of keto-adaptation is significantly lower in KD-fed rats that transitioned from TRF (red) relative to *ad libitum*-fed animals (black). However, rats that transitioned from SD to KD with a previous cycle of KD (blue) showed the lowest GKI. (C) Consumption of the SD was associated with high baseline GKI levels immediately after the cessation of carbohydrate restriction. Data are represented as group mean ±1 SEM, * p < 0.05.

As in the previous cohorts of rats, when dietary ketosis with TRF was initiated following *ad libitum* feeding, aged rats showed significant delays in keto-adaptation (F_[1,21]_ = 21.05; p < 0.001). Keto-adaptation was compared between KD-naïve rats that cycled into ketosis following *ad libitum* feeding (Group A cycle 1) to rats that were KD-naïve following 4 weeks of TRF (Group B cycle 2). Overall, across days 1-7 on the KD, the GKI significantly decreased as a function of days on the diet (F_[2,28]_ = 22.89; p < 0.001). The GKI, however, was significantly lower in Group B during cycle 2, compared to Group A during cycle 1 (F_[1,14]_ = 16.74; p = 0.001), indicating that 4 weeks of fasting through TRF facilitated the ability to flexibly switch from glycolysis to ketosis compared to consuming food *ad libitum*, presumably by decreasing insulin levels. Across both cycles of ketosis initiation, the significant main effect of age persisted (F_[1,14]_ = 10.44; p = 0.001), with aged rats having higher GKI values relative to young animals. Interestingly, there was a significant interaction effect between age and cycle (F_[1,14]_ = 8.20; p = 0.01), indicating that the fasting through TRF expedited keto-adaptation in aged rats to a greater extent than young. This interaction is likely due to metabolic impairments and hyperinsulinemia of aged rats while being fed *ad libitum*, which was at least partially being overcome by TRF.

To examine the impact of previous experience on the KD to the rate of keto-adaptation in young and aged animals, we then compared GKI values from rats that were naïve to dietary intervention (Group A cycle 1) to their GKI values during their second cycle of ketosis (Group A cycle 3). GKI levels were significantly lower during the second keto-adaptation cycle than the first (F_[1,14]_ = 52.12; p < 0.001). Moreover, cycle significantly interacted with age (F_[1,14]_ = 6.66; p = 0.02), indicating that the age-associated delay in keto-adaptation was mitigated by previous experience on the KD.

Because TRF with the SD was sufficient to influence keto-adaptation, the degree to which previous experience in ketosis influenced successive cycles of keto-adaptation was also examined. Specifically, Group A cycle 3 (TRF with prior KD experience) was compared to Group B cycle 2 (TRF and KD naïve). Critically, both groups had 1 month of TRF prior to transitioning to the KD, with these groups differing only in terms of previous experience on the KD. Importantly, the GKI was significantly lower in Group A cycle 3 compared to Group B cycle 2 (F_[1,14]_ = 15.89; p = 0.001). This observation indicates that a previous cycle of keto-adaptation enhanced the rate of metabolic switching to promote higher rates of ketogenesis in a shorter amount of time following carbohydrate restriction. In other words, the increased rate of keto-adaptation induced by 4 weeks of TRF was further augmented by previous experience with keto-adaptation. In contrast to Cycle 1 in Group A (first cycle of keto-adaptation following *ad libitum* feeding), there were no significant effects of age (F_[1,14]_ = 2.43; p = 0.14) across the two later cycles.

### Keto-adaptation in young and aged rats is decoupled from glycogen content

One hypothesis regarding the initiation of ketosis is that fat oxidation is promoted when glycogen content in the liver and muscles is depleted. The age-related delay in keto-adaptation observed in aged animals could therefore be due to increased glycogen levels in liver or muscle. Thus, to determine how glycogen dynamics interact with ketosis, we measured the glycogen content in the liver and muscle of young and aged rats on the SD or KD for at least 12 weeks prior to sacrifice. Most rats were fasted for at least 15 hours prior to sacrifice, however a small subset of rats was fed 1 hour prior to sacrifice. While there were no differences in liver glycogen content between age groups (F_[1,50]_ = 2.50; p = 0.12; Figure 8A), there was a significant decrease in liver glycogen in KD-fed rats (F_[1,50]_ = 6.60; p = 0.01). Additionally, for the SD-fed rats only, fasted rats had significantly less glycogen in their liver than fed rats (F_[1,50]_ = 4.88; p = 0.03). This difference between fasted and unfasted animals was particularly evident in the aged rats and was not observed in the KD group, indicating that ketosis prevents fasting-induced glycogen depletion. These data suggest that long-term ketosis reduced liver glycogen content, but these levels were not affected while fasting. In contrast, aged rats on a SD that were fasted quickly lost liver glycogen, which was not observed in young rats. There were no differences in glycogen content within the muscle across age (F_[1,50]_ = 0.47; p = 0.50), diet (F_[1,50]_ = 0.04; p = 0.85) or fasted versus fed groups (F_[1,50]_ = 1.31; p = 0.26; Figure 8B). Furthermore, there were no significant interactions between any of these variables (p > 0.14 for all).

**Figure 8:**
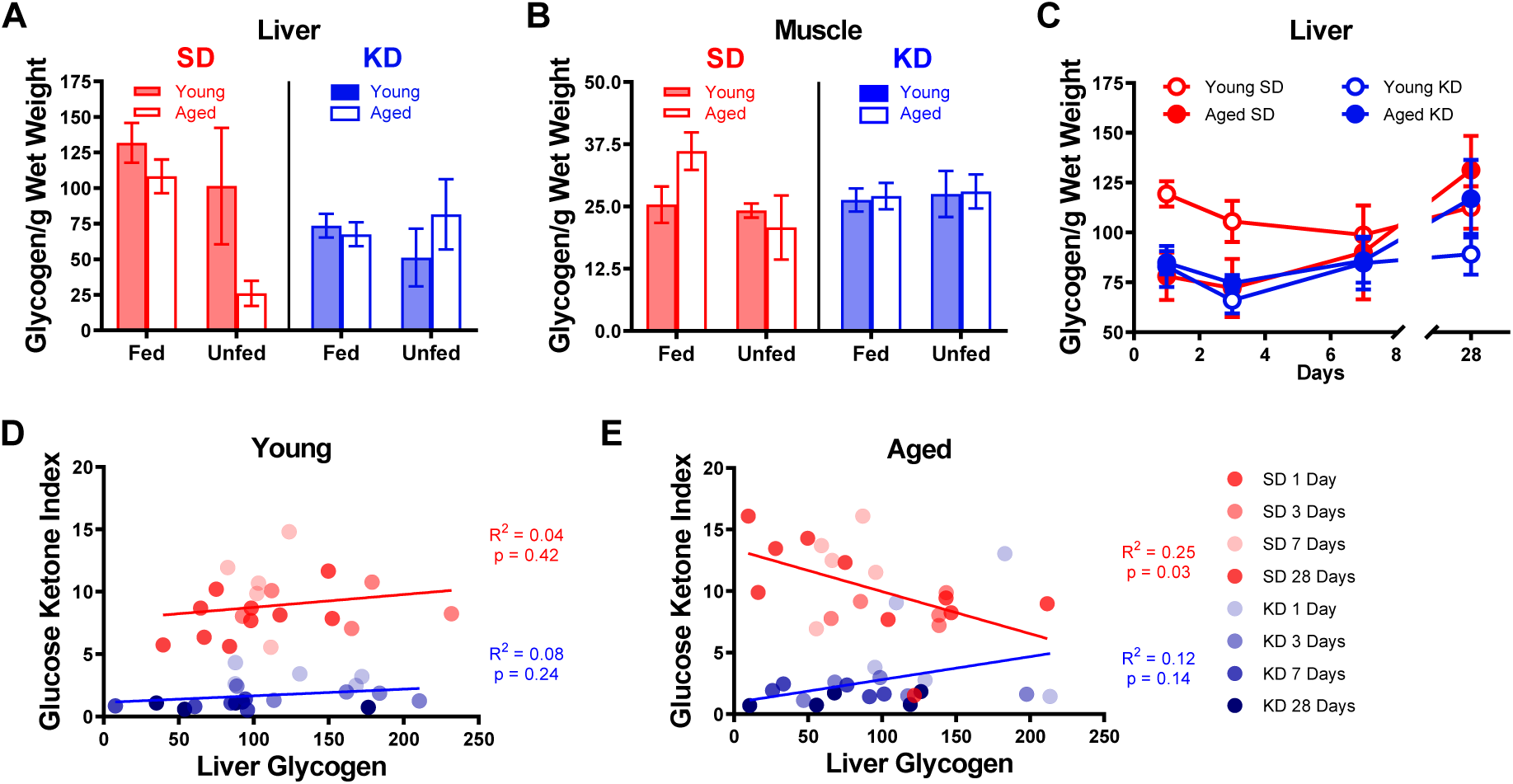
Keto-adaptation does not rely on glycogen depletion. (A) Glycogen levels in the liver while fasted or fed in young and aged rats on the SD or KD. SD-fed aged rats were not able to maintain liver glycogen when fasted, whereas KD-fed aged rats were. (B) Muscle glycogen while fasted or fed in young and aged rats on the SD or KD did not change across age, diet or feeding status groups. (C) Young SD-fed rats had significantly higher liver glycogen than all aged rats and KD-fed young rats during the first 3 days of the diet, but levels were similar in all groups on day 7 and beyond. (D) The GKI did not correlate with the amount of glycogen in the liver during keto-adaptation for young animals on the SD or KD. (E) The GKI did not correlate with the amount of glycogen in the liver during keto-adaptation for aged animals on the KD, but did significantly negatively correlate during this time period in SD-fed rats. Data are represented as group mean ±1 SEM, * = p < 0.05.

To determine if the glycogen content within the liver changed during the process of keto-adaptation, 40 young and 40 aged rats were split into groups 1-4, which were fed 51 kcal once daily of the KD or SD for 1, 3, 7 or 28 days and then sacrificed. Because age differences in liver glycogen were only detected in fasted animals, all rats were sacrificed 18-22 hours after eating for quantification of liver glycogen content. Consistent with our previous data, following 1 and 3 days of consuming the SD, liver glycogen was significantly lower in aged rats relative to young (t_[8]_ = 3.00; p = 0.02 and t_[8]_ = 3.23; p = 0.01, respectively). In the KD group at days 1 and 3, there was no age difference in liver glycogen (t_[8]_ = 0.51; p = 0.62 and t_[8]_ = 0.16; p = 0.88, respectively). By days 7 and 28, there was no longer a significant difference in liver glycogen content across age or diet groups nor did these two factors significantly interact (p > 0.26 for all comparisons). When time point was considered as a factor, there was a significant change in glycogen levels by day (F_[3,54]_ = 3.18; p = 0.03), as well by diet (F_[1,54]_ = 6.01; p = 0.02). While there was no significant effect of age group (F_[1,54]_ = 1.8; p = 0.17), this factor did significantly interact with length of time on the diet (F_[1,54]_ = 5.26; p = 0.03). This could be due to the tendency for glycogen levels to increase after 3 days on the diet in KD-fed rats.

Surprisingly, in young rats, liver glycogen content did not correlate with the GKI during the initiation of TRF in either KD-fed rats (R^2^ = 0.08; p = 0.24) or SD-fed rats (R^2^ = 0.04; p = 0.42; Figure 8D). Furthermore, there was no correlation between liver glycogen and GKI in aged KD-fed rats during the initiation of ketosis (R^2^ = 0.12; p = 0.14; Figure 8E). However, in the SD-fed rats that switched from *ad libitum* feeding to TRF, liver glycogen in aged SD-fed rats significantly negatively correlated with GKI (R^2^ = 0.25; p = 0.03). These data demonstrate that it is not critical for liver glycogen to be depleted to initiate dietary ketosis in either aged or young animals. Moreover, in aged rats that are transitioning from *ad libitum* feeding to TRF of a SD, more glycogen is associated with lower GKI values. This suggests that the aged rats that adapt to being fasted by decreasing blood glucose and increasing BHB level spare liver glycogen content.

### Metabolic responses to a physiological stressor are enhanced by ketosis, but not time-restricted feeding

To investigate whether liver glycogen could still be utilized in ketotic animals, the metabolic response to stress, which involves a rapid hepatic conversion of glycogen to glucose (Sherwin and Saccà, 1984), was pharmacologically induced. Importantly, this evolutionarily conserved metabolic response to a stressor can be essential for properly adapting to activities of daily living, especially during strenuous circumstances. To investigate this, serum levels of glucose and BHB following epinephrine injections were quantified in rats fed *ad libitum* with traditional rodent chow (Envigo, Teklad 2918). These same animals were then placed on the KD or SD for 12 weeks. At both time points, rats were given an intraperitoneal injection of either saline or 0.1mg/kg epinephrine 30 minutes postprandially, and blood glucose and BHB levels were quantified 30 minutes post-injection. Injections were separated by a 48-hour washout period, with the order counterbalanced across animals. The post-injection values of blood glucose, BHB and the GKI for saline and epinephrine were then used to calculate the percent change between saline and epinephrine injections for blood levels of glucose (Figure 9A), BHB (Figure 9B) and the GKI (Figure 9C).

**Figure 9:**
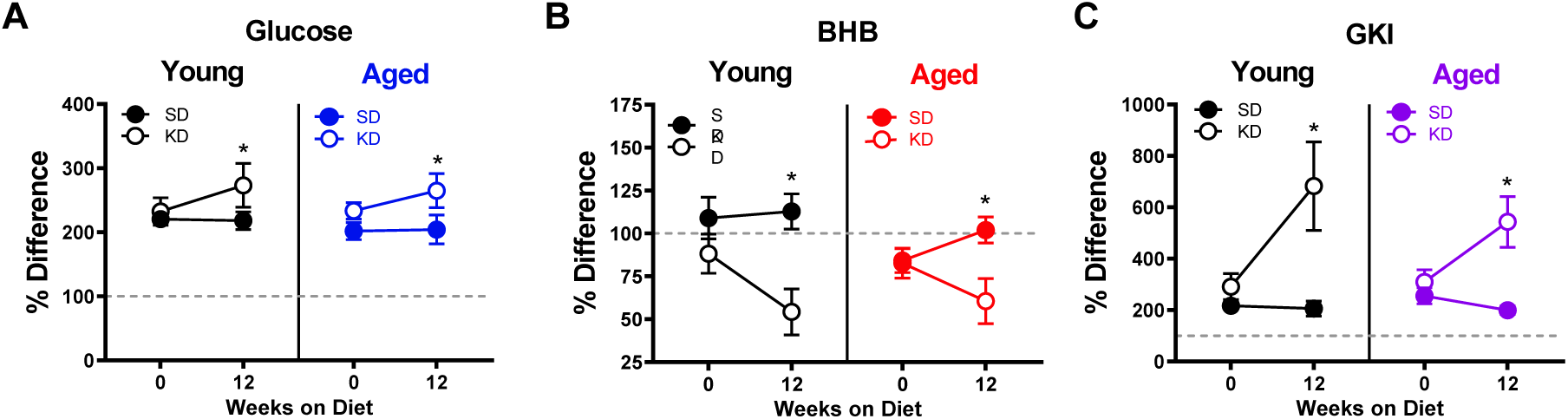
Keto-adaptation enhances physiological metabolic response to stress. The percent difference in blood (A) glucose, (B) BHB and (C) GKI between saline and epinephrine injections during *ad libitum* feeding and TRF with SD or KD. Data are represented as group mean ±1 SEM, * = p < 0.05.

All rats, including those on the KD, showed a marked increase in blood glucose in response to epinephrine. In fact, there was a significant effect of diet on epinephrine-induced increases in blood glucose, with KD-fed rats having a larger change relative to the baseline saline condition (F_[1,23]_ = 5.21; p = 0.032). This observation indicates a conservation of the stress-induced glucose response while in ketosis. The enhanced ability to mobilize glucose in response to a stressor was not due to the TRF alone, as the SD-fed rats did not show this same enhancement between *ad libitum* feeding and 12 weeks of the SD (F_[1,13]_ = 0.29; p = 0.599).

In line with these data, blood levels of the pancreatic hormone glucagon, which is required for the breakdown of glycogen to glucose, was not altered in KD-fed rats (F_[1,13]_ = 0.77; p = 0.394; Figure 10A). While we did not observe an age difference in our SD-fed rats, when older groups of animals were included there was a significant difference between young and old animals (Figure 10B-C), replicating previously published work (Gold, 2005; Korol, 2002).

**Figure 10:**
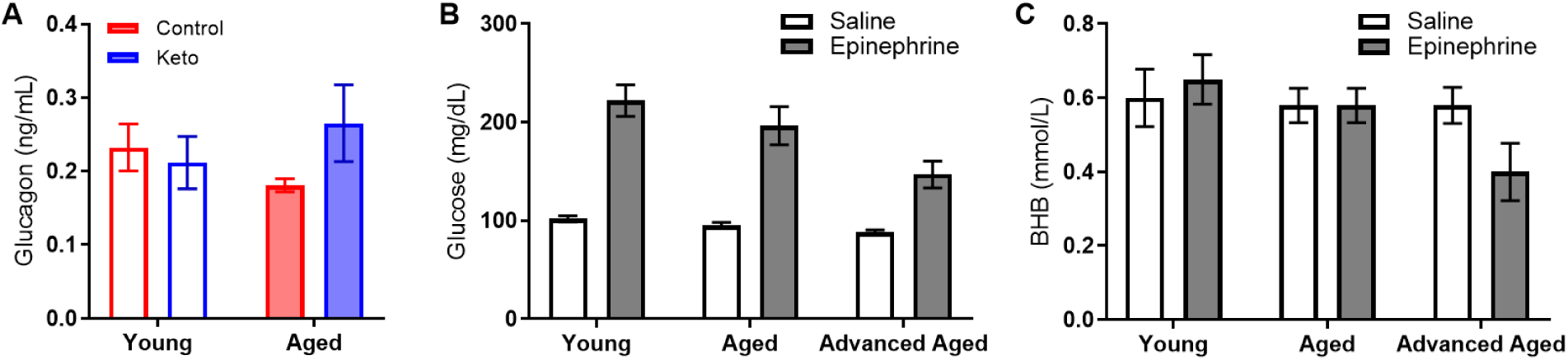
Glucagon, glucose and BHB response to epinephrine change as a function of advancing age. In rats of this strain there is not an age-related impairment in the glucose response to epinephrine at 24 months, but there is as 33 months in both (B) glucose and (C) BHB following epinephrine injections. Data are represented as group mean ±1 SEM, * = p < 0.05.

Although BHB levels were unaffected by epinephrine during *ad libitum* feeding with chow or during TRF with the SD, epinephrine decreased blood BHB levels in KD-fed rats. Moreover, in rats that went from *ad libitum* feeding to TRF with the KD, there was a significant change in the percent decrease in BHB induced by epinephrine (F_[1,23]_ = 8.90; p = 0.007). In KD-fed rats, the glucose increase and concurrent decrease in BHB, was associated with a significant epinephrine-induced increase in GKI that did not occur during *ad libitum* feeding (F_[1,23]_ = 15.39; p = 0.001). Despite the isocaloric 51 kcal per day across both diet groups, the abrupt increase in the post-epinephrine GKI experienced by the KD-fed rats was not observed in the SD-fed group, indicating that TRF alone did not enhance the metabolic response to stress in either young or aged rats. The present observations suggest that long-term nutritional ketosis does not deplete glycogen stores. Moreover, hepatic glycogen can be mobilized in response to a physiological stressor to a greater extent when in ketosis relative to glycolysis alone.

## DISCUSSION

Here we show that age-related metabolic dysfunction is mitigated by changing the energy consumption pattern to time-restricted feeding (TRF; once daily feeding) of 51 kcal. These benefits in metabolic health were further bolstered by a high fat, low carbohydrate ketogenic diet (KD). We replicated previous data (Hernandez et al., 2018; Hernandez et al., 2018a) demonstrating an age-related impairment in keto-adaptation rate, and extended these observations to show that keto-adaptation induced by diet does not correlate with liver glycogen depletion. Surprisingly, when fasted, rats in long-term ketosis had levels of liver and muscle glycogen that were comparable to young rats on the standard diet (SD) and considerably higher than aged SD-fed rats. Moreover, the KD appeared to enhance glycogen dynamics, with KD-fed rats showing an enhanced increase in blood glucose in response to epinephrine. Thus, age-related declines in metabolic switching do not appear to result from differences in glycogen storage and dynamics, but rather emerge from metabolic dysfunction associated with hyperinsulinemia.

Aged rats had elevated fasting and postprandial insulin levels while *ad libitum* feeding with standard rodent chow, as well as during the initial stages of keto-adaptation. In young rats, fasting insulin levels decreased significantly following the first 24 hours of a KD. This decrease in fasting insulin was not observed in aged rats during the first week of a KD. In addition to insulin deficits, aged animals fed *ad libitum* with chow had significantly elevated leptin, c-peptide, and HOMA. While some metabolic factors were significantly improved following TRF with the SD, such as the insulin tolerance test, combining TRF with a KD was most efficacious at restoring metabolic markers in aged rats to levels comparable to young animals, as well as enhancing the epinephrine-induced increase in peripheral glucose.

If metabolic switching is defined as being able to switch bi-directionally between periods of carbohydrate/glucose usage and fatty acid/ketone body usage (Mattson et al., 2018), then the KD enhanced metabolic switching beyond what was observed with TRF alone. While TRF did increase the rate at which aged rats could transition from glycolysis to ketosis, enhanced keto-adaptation was further augmented by TRF+KD. These data concur with previous studies demonstrating the additive benefits of food restriction with a KD at increasing seizure threshold (Bough et al., 1999). Surprisingly, young and aged rats in our study consuming the KD still showed an enhanced glucose response to an acute increase in epinephrine, beyond what was observed in rats fed TRF+SD. This observation further supports the notion that TRF+KD may confer the greatest improvement in metabolic switching. Enhancing metabolic switching may be particularly important in aged populations. While epinephrine is able to improve memory performance in young rats through hepatic glucogenesis, aged rats show an attenuated elevation in blood glucose in response to epinephrine, and no cognitive benefits. Thus, old animals do not cognitively benefit from this physiological stress response that is an evolutionary conserved mechanism to promote survival (Gold and Korol, 2014).

Although TRF alone can enhance metabolic switching, and improve cognitive health across the lifespan (Anson et al., 2003; Harvie et al., 2013, 2011; Hatori et al., 2012; Mattson et al., 2018; Mitchell et al., 2019), the current data suggest that additional benefits can be obtained through nutritional ketosis. It is also notable that the benefits of TRF for cognitive function might be due to comparisons with *ad libitum* fed rodents that are metabolically morbid (Martin et al., 2010). In fact, it is well documented that excessive energy consumption can lead to cognitive impairments (Bocarsly et al., 2015; Kanoski and Davidson, 2011; Stranahan et al., 2008). A strong indicator that TRF alone is not sufficient for extending the cognitive healthspan is the wealth of studies demonstrating age-related cognitive decline in rodents while utilizing tasks requiring appetitive motivation. Many tests of cognitive function require animals to be restricted to <85% of their body weight to motivate participation by feeding once daily, making a large portion of investigations on the neurobiology of cognitive aging into incidental experiments of caloric restriction with TRF. While incentive motivation for food rewards does not change with age (Simon et al., 2010), robust age-related deficits are observed on working memory (Bañuelos et al., 2014; Beas et al., 2016; Caesar M. Hernandez et al., 2018), other executive functions (Barense, 2002; Hernandez et al., 2017; Samson et al., 2014), as well as behaviors that require the prefrontal cortex and medial temporal lobe (Hernandez, Hernandez, Campos, Truckenbrod, Federico, et al., 2018; Hernandez, Reasor, Truckenbrod, Campos, Federico, et al., 2018; Hernandez et al., 2015; Johnson et al., 2017).

Importantly, there may be pertinent differences in metabolism, brain aging, and responses to dietary ketosis between males and females (Zhao et al., 2016). Unfortunately, the current lack of availability of female Fischer 344 × Brown Norway hybrid rats from maintained rodent colonies (https://www.nia.nih.gov/research/dab/aged-rodent-colonies-handbook) did not permit the consideration of sex as a variable in the current experiments. Potential interactions of sex and dietary ketosis will need to be examined in future studies.

We’ve shown that combining TRF with a KD provides a more robust restoration of metabolic health than TRF alone. In addition to the metabolic factors measured here, a previous study has shown that this feeding pattern can enhance cognitive function and decrease anxiety-like behaviors (Hernandez et al., 2018a). Moreover, TRF+KD alters transporter protein expression within the prefrontal cortex and hippocampus (Hernandez et al., 2018a; Hernandez et al., 2018), increases BDNF (Garriga-Canut et al., 2006; Masino and Rho, 2012), decreases pain and inflammation (Ruskin et al., 2009), modulates K_ATP_ channels (Ma et al., 2007; Masino and Rho, 2012), modifies neurotransmitter levels (Yudkoff et al., 2008, 2005) and lessens amyloid and tau pathologies (Kashiwaya et al., 2013). Each of these biochemical alterations induced by ketosis are likely to confer resilience to cognitive decline.

In addition to peripheral health, memory deficits and other types of cognitive impairment have a profound contribution to decreasing quality of life in individuals and their caregivers. Although the increased mean life expectancy has had a commensurate increase in the number of aged people living without physical limitations (Christensen et al., 2009), the same cannot be said for cognitive deficits. In fact, the proportion of individuals experiencing mild or severe memory impairments will reach 1 in 3 by the 8^th^ decade of life (Blazer et al., 2015), and disrupted brain bioenergetics is a common feature of aging (Cunnane et al., 2011; Mattson and Arumugam, 2018). Improving metabolic function in the aged population can have profound impacts on cognitive outcomes, as the brain disproportionately uses metabolic resources (Cunnane et al., 2011) Furthermore, cerebral glucose metabolism decreases with age, most notably within the hippocampus and prefrontal cortex (Gage et al., 1984), both of which are critical for higher cognitive function (Frith and Dolan, 1996; Morris et al., 1982; Sloan et al., 2006). Importantly, this decline correlates with worse spatial memory performance (Gage et al., 1984). Thus, developing metabolic-based interventions for improving peripheral metabolic health will likely have profound therapeutic potential for increasing the cognitive healthspan.

## Materials and Methods

### Subjects & Handling

Young (4 months; n = 112; mean weight 337.59g) and aged (20 months: n = 112; mean weight 526.04) male Fischer 344 × Brown Norway F1 Hybrid rats from the NIA colony at Charles River were used in this study. Rats were housed individually and maintained on a reverse 12-hr light/dark cycle, and all feeding, metabolic and biochemical assays were performed in the dark phase. Rats were given one week to acclimate to the facility prior to any intervention. All experimental procedures were performed in accordance with National Institutes of Health guidelines and were approved by the Institutional Animal Care and Use Commity at the University of Florida.

### Dietary interventions

For all experiments, the same ketogenic diet (KD) and standard diet (SD) were used as published previously (Hernandez et al., 2018a; Hernandez et al., 2018; Hernandez et al., 2018b). An additional group of rats were fed *ad libitum* with standard laboratory chow (Envigo, Teklad 2918). The KD was a high fat/low carbohydrate diet (Lab Supply; 5722, Fort Worth, Texas) mixed with MCT oil (Neobee 895, Stephan, Northfield, Illinois) with a macronutrient profile of 76% fat, 4% carbohydrates, and 20% protein (see Figure 1A). The micronutrient-matched SD (Lab Supply; 1810727, Fort Worth, Texas) had a macronutrient profile of 16% fat, 65% carbohydrates, and 19% protein (see Figure 1A). The KD and SD were made fresh weekly and kept at 4°C until use each day. For additional micronutrient details for both diets, see Hernandez et al., 2018a. The Teklad standard laboratory chow had a macronutrient composition of 5% fat, 75% carbohydrates and 20% protein. The SD and KD were micronutrient matched, and rats were fed the same number of calories each day (∼51kcal) during the dark phase between 14:00 and 17:00 hours (lights on at 20:00) while on TRF, regardless of diet. Access to water was *ad libitum* for all rats on all diets, and rats were weighed daily when fed. These different diets are reported to not alter water consumption patterns or induce a prolonged stress response (Hernandez et al., 2018a).

### Blood collection, testing, and metabolic data

All blood samples collected prior to sacrifice were collected from a small nick in the tail. Rats were briefly restrained, tails were cleaned with an alcohol swab, and then a clean scalpel blade was used to make a superficial cut approximately 1 inch from the end of the tail. Following blood collection, wounds were wiped clean and Wonder Dust (Farnam Companies, Phoenix, AZ) was applied to promote wound healing. Blood collected on the day of sacrifice was collected directly from the trunk immediately following decapitation.

For blood glucose and ketone body (β-hydroxybutyrate; BHB) determination, one drop per test was collected directly onto the appropriate test strip (Abbott Diabetes Care, Inc, Alameda, CA; glucose SKU#: 9972865 and ketone SKU#: 7074565) in a Precision Xtra blood monitoring system (Abbott Diabetes Care, Inc, Alameda, CA; SKU#: 9881465).

For the insulin tolerance test (ITT) and glucose tolerance test (GTT), rats were fasted overnight and injected intraperitoneally with either 0.75 U/kg of human insulin (Hanna Pharmaceutical Supply Co #NC0769896) or 1 g/kg glucose (Fisher Scientific #D16-500). Blood glucose and ketone body levels were measured as described above at 0, 15, 30, 45, 60, and 120 minutes post injection.

For metabolic biomarker quantification, approximately 750 μL of blood was collected from the tail nick at the time of glucose testing. Collected blood was left at room temperature for at least 25 minutes prior to centrifugation at 13,000g for 10 minutes at 4°C. 0.35 μL of supernatant was collected into a tube containing 0.4 μL DPP-IV inhibitor (EMD Millipore, NC9010908), 0.4 μL protease inhibitor cocktail (Millipore Sigma, S8820-2TAB) and 0.32 μL 0.9% PBS. Samples were kept frozen at −80°C prior to use in the Rat Metabolic Array (MRDMET; EVE Technologies, Calgary, AB). This multiplex immunoassay (BioPlex 200) quantified levels of insulin, leptin, C-peptide 2 and glucagon. Each sample was analyzed in duplicate. All values outside of the liner range were excluded.

### Epinephrine Administration

Prior to starting any dietary intervention, half of the rats were given 0.1mg/kg epinephrine (diluted in 0.9% sterile saline solution) and half were given the equivalent volume of saline delivered via intraperitoneal injection. After 30 minutes, glucose and BHB were determined as described above. Groups were counterbalanced for drug across two days of testing with one day off in between. That is, following a 48-hour washout period, the groups were reversed and the process was repeated. This entire 2-day process was repeated after 12 weeks on the TRF+SD and TRF+KD fed rats, with epinephrine injections occurring 30 minutes postprandially.

### Tissue Collection and Glycogen Quantification

On the day of sacrifice, a subset of rats was fed their respective diets one hour prior to being anesthetized. All other rats were sacrificed without feeding since the prior day (approximately 15-22 hours fasted). All rats were rapidly sedated in a jar containing concentrated isoflurane and then decapitated once the righting reflex had ceased. The liver and tibialis anterior muscle were isolated, rinsed in cold PBS and immediately frozen on dry ice. Tissue was stored at −80°C until use.

Glycogen content in the liver and muscle tissue were determined per mg wet tissue weight, modeled after previously published protocols (Passonneau et al., 1967; Passonneau and Lauderdale, 1974; Shetty et al., 2012). Approximately 50 mg frozen tissue was placed in a pre-weighed tube containing 660 μL NaOH at 100°C and tissue weight was determined. Tubes were boiled at 100°C for 15 minutes, vortexed and then brought to neutral pH using 320 μL 2M HCl. Samples were vortexed again and 150 μL of muscle, or 20 μL of liver with 130 μL of ddH_2_O, were placed into two tubes each. A sodium acetate solution was added to each tube, one with and one without the addition of amyloglucosidase, and samples were incubated at 37°C for 2 hours. Following a 20-minute centrifuge at 14,000 RPM, 50 μL of supernatant was plated in triplicate into a 96 well plate. 100 μL Tris buffer was added to each well, which was then read on a Synergy HT Microplate Spectrophotometer (BioTek) at 340 nm. 50 μL hexokinase solution was added to each well, and the plate was incubated at room temperature on a shaker, away from light before reading again at 340 nm. The optical density reading before the addition of hexokinase was subtracted from the final optical density reading to correct for free glucose values. Final glycogen content was extrapolated using a standard glycogen curve run on each plate and normalized to tissue weight.

### Quantification and Statistical analysis

All data are expressed as group means ± standard error of the mean (SEM) unless otherwise reported. The glucose ketone index (GKI) was calculated as described in a previous publication (Meidenbauer et al., 2015). Briefly, the glucose (mg/dL) was divided by the molecular weight of glucose (expressed as (mol/(g × dL))), and this value was then divided by the BHB value (mM). Differences across age and diet groups were analyzed using a two-factor ANOVA with the between subjects factors of age (2 levels: young and aged) and diet (2 levels: SD and KD or 3 elevels: SD, KD and *ad libitum*). When time was a factor, an ANOVA-RM was used with the same factors. Null hypotheses were rejected at the level of p > 0.05. All analyses were performed with the Statistical Package for the Social Sciences v25 (IBM, Armonk, NY) GraphPad Prism version 7.03 for Windows (GraphPad Software, La Jolla, California USA).

## Supplementary Material

Additionally, a glucose tolerance test (GTT) was administered during the 7^th^ week of dietary intervention. While the glucose injection did result in increased circulating glucose expression over time (F_[6,186]_ = 34.09; p < 0.001; Figure S1A), there were no differences across age (F_[1,31]_ = 0.14; p = 0.71) or diet groups (F_[2,31]_ = 1,15; p = 0.33), and none of these variables significantly interacted (p ≥ 0.10 for all comparisons). Furthermore, there were no differences across age (F_[5,1]_ = 0.28; p = 0.60) or diet (F_[5,2]_ = 2.76; p = 0.08) in the AUC (Figure S1B). However, injecting glucose did alter circulating levels of BHB differentially across diet groups (F_[2,31]_ = 17.30; p < 0.001), and diet group significantly interacted with the amount of time post injection (F_[12,186]_ = 4.55; p < 0.001; Figure S1C). While there was a significant effect of time point for all groups (F_[6,186]_ = 31.08; p < 0.001), there were no differences in BHB response across age groups (F_[1,31]_ = 17.30; p = 0.06) nor did age interact with time point (F_[6,186]_ = 1.54; p = 0.17). However, age did significantly interact with diet group (F_[1,31]_ = 3.46; p = 0.04) such that there was a strong trend for aged free-fed rats to demonstrate lower levels than their diet-matched young counterparts (F_[1,8]_ = 4.91; p = 0.06), whereas aged KD-fed (F_[1,11]_ = 3.25; p = 0.10) and aged SD-fed rats (F_[1,12]_ = 1.31; p = 0.28) did not differ from their diet-matched young counterparts. Finally, there were no differences across age (F_[5,1]_ = 0.25; p = 0.62) or diet (F_[5,2]_ = 0.63; p = 0.54) in the AUC (Figure S1D).

**Figure S1:**
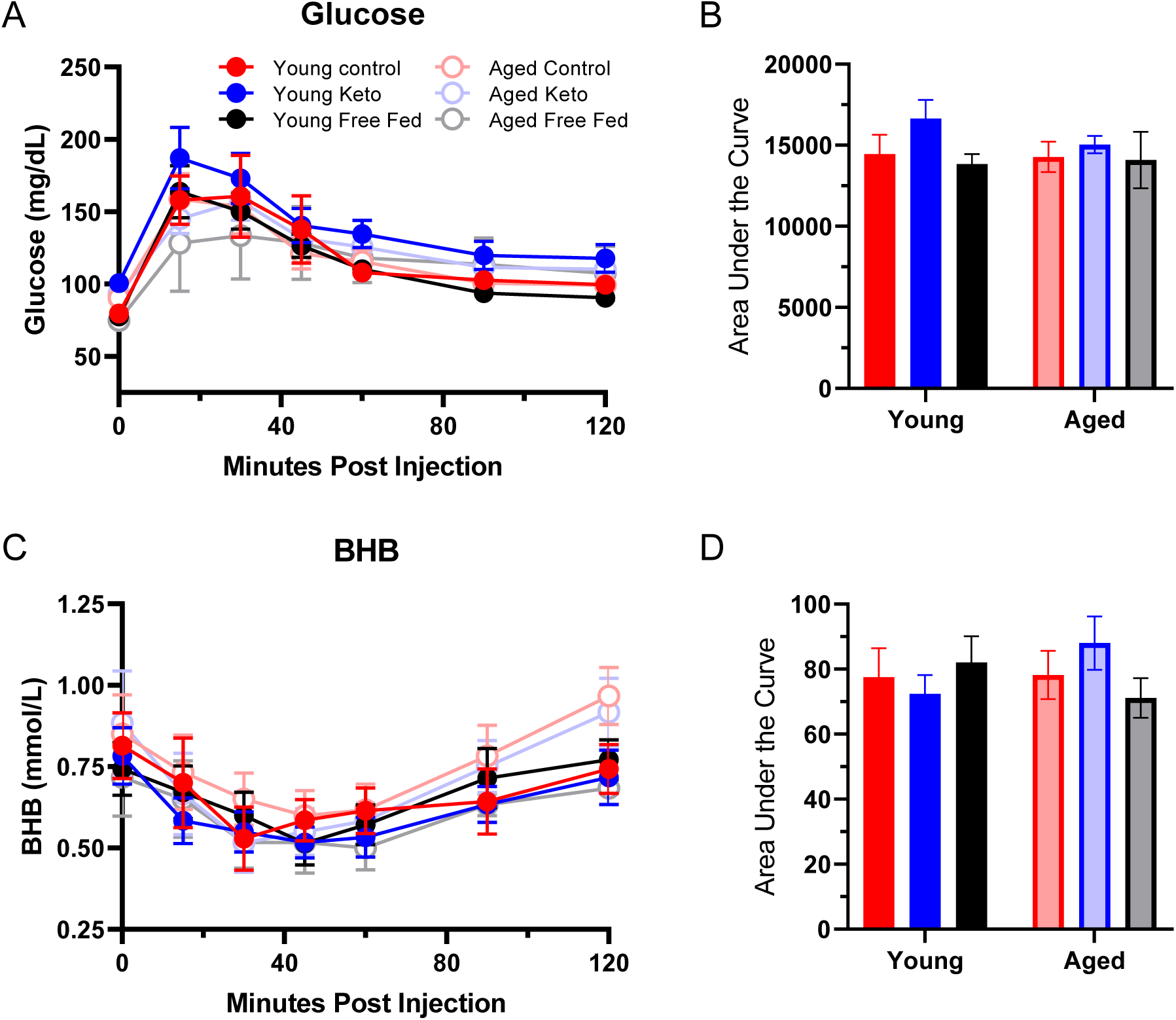
Glucose tolerance testing (GTT) following 7 weeks dietary implementation. (A) There were no differences in (A) glucose values over time nor (B) area under the curve (AUC) across age or diet groups following an injection of a bolus of glucose intraperitoneally. However, diet did significantly affect (C) BHB values over time, (D) but no differences across age or diet groups were observed in the total AUC following glucose injection. Data are represented as group mean ±1 SEM.

